# Small but significant genetic differentiation among populations of *Phyllachora maydis* in the midwestern United States revealed by microsatellite (SSR) markers

**DOI:** 10.1101/2023.10.31.563566

**Authors:** Tiffanna. J. Ross, Blaise Jumbam, John Bonkowski, Jennifer L. Chaky, Martin I. Chilvers, Stephen B. Goodwin, Nathan. M. Kleczewski, Daren. S. Mueller, Alison. E. Robertson, Damon. L. Smith, Darcy. E. P. Telenko

**Affiliations:** Department of Botany and Plant Pathology, Purdue University, West Lafayette, IN 47907, U.S.A; Corteva Agriscience, Johnston, IA, 50131, U.S.A; Department of Plant, Soil and Microbial Sciences, Michigan State University, East Lansing, MI 48824, U.S.A; Crop Production and Pest Control Research Unit, U.S. Department of Agriculture-Agricultural Research Service (USDA-ARS), Purdue University, West Lafayette, IN 47907, U.S.A; Department of Crop Sciences, University of Illinois, Urbana, IL 61801, U.S.A; Department of Plant Pathology and Microbiology, Iowa State University, Ames, IA 50011, U.S.A; Department of Plant Pathology, University of Wisconsin-Madison, Madison, WI 53706, U.S.A

**Keywords:** corn, tar spot, genetic diversity, gene flow, mode of reproduction, molecular markers

## Abstract

*Phyllachora maydis* Maubl, the causal pathogen of tar spot of corn (*Zea mays* L.), has emerged recently in the United States and Canada. Studies related to its genetic diversity and population structure are limited and are necessary to improve our understanding of this pathogen’s biology, ecology, epidemiology, and evolutionary potential within this region. This study developed and used 13 microsatellites (SSR markers) to assess the genetic population structure, diversity, gene flow and reproductive mode of 181 *P. maydis* samples across five states in the Midwest U.S. The polymorphic information content (PIC) of loci ranged from 0.32 to 0.72 per locus, indicating their high utility for assessing the dynamics of *P. maydis* populations. Analysis of molecular variance (AMOVA) detected a significantly low, but statistically significant genetic differentiation (F_ST_ = 0.15) among populations, where 85% of the variance resided within populations. *P. maydis* populations were highly diverse (He = 0.55), with moderate gene flow (Nm = 2.80), and showed evidence of sexual recombination (*r̄_<u>d</u>_*; *p = > 0.001*). Structure analysis showed the samples were not geographically structured but rather grouped into two genetic clusters (k =2) of severe genetic admixture suggesting possible long-distance dispersal of aerial spores or infected corn materials among the five Midwest states. Both principal coordinate analysis (PCoA) and discriminate analysis of principal component (DAPC) supported the STRUCTURE analysis of the two clusters. These 13 highly polymorphic molecular markers could be used for future investigations of this pathogen’s population dynamics within the U.S., and possibly populations outside.

## Introduction

Corn (*Zea mays* L.) is the largest staple food crop produced globally, at 1.2 billion tons per year (USDA-NASS 2021). It is an essential source of animal feed, food, fuel, and export in the United States (USDA-NASS 2021) but is susceptible to several diseases incited by many plant pathogens (Mueller et al. 2020; Wise et al. 2016). In 2015, a new foliar disease of corn, tar spot, was detected in the U.S. in Illinois and Indiana but without significant economic impact (Ruhl et al. 2016). However, in 2018, several corn-producing areas in the Midwest experienced their first major epidemic of tar spot in which 4.7 million metric tons (183.6 million bushels) worth $US 658.7 million were lost (Crop Protection Network, 2023; Mueller et al. 2020). Tar spot was later detected in Ontario, Canada in 2020 (Tenuta 2020). Tar spot has become one of the most significant and severe diseases of corn in the Midwest and Ontario, Canada in terms of economic and yield losses in recent years (Mottaleb et al. 2019; Mueller et al. 2022). In 2021, 5.97 million metric tons (235.2 million bushels) of corn yield were lost in the U.S. and Ontario, Canada valued at $US 1,268 million (Crop Protection Network, 2023; Mueller et al. 2022).

Tar spot of corn is caused by *Phyllachora maydis* Maubl, an obligately biotrophic fungal pathogen belonging to the order *Phyllachorales* in the class Sordariomycetes (Bajet et al. 1994; Maublanc 1904; McCoy et al. 2019; Parbery 1963). This fungus is most likely haploid and is thought to have a heterothallic mating system (MacCready et al., 2023). The *Phyllachorales* contains approximately 160,000 species known globally, of which 1,226 are currently acknowledged (Cannon 1997; Kirk et al. 2008; Maharachchikumbura et al. 2016; Mardones et al. 2017). Broders et al. (2022) and Mardones et al. (2017) conducted a comprehensive assessment of *P. maydis*, providing evidence that understanding of this species and its genera is limited and requires significant attention. Species found in the order *Phyllachorales* are associated with many graminaceous plants but can also infect dicots. They are presumed to be highly host specific but are found on hosts across a vast range of habitats. Although the assumptions of host specificity do not always hold for some genera within the *Phyllachorales*, so far corn is the only known host for *P. maydis* (Cannon 1991 and 1997; Cline 2019; Kleczewski et al. 2020; Parbery 1963; Valle-Torres et al. 2020).

*Phyllachora maydis* was first identified in Mexico during 1904 and was detected in 18 other countries in the Caribbean, Central, and South America during the past century (Valle-Torres et al. 2020). Since the first detection of *P. maydis* in the U.S. during 2015, it has spread to at least 14 states and to Ontario, Canada, and its range continues to expand every year (https://corn.ipmpipe.org/tarspot/; Athey 2020; Collins et al. 2021; Dalla Lana et al. 2019; Malvick et al. 2020; McCoy et al. 2018; Pandey et al. 2022; Ruhl et al. 2016; Tenuta 2020). *Phyllachora maydis* has two reproductive states; the sexual state produces ascospores and the asexual, conidia. Infection is presumed to occur primarily via the ascospores, which are thought to overwinter in asci in perithecia produced in stromata on corn residue (Groves et al. 2020; Kleczewski et al. 2019). Spore dissemination up to 1,200 m from the inoculum source by wind or rain to nearby susceptible plants has been documented (Kleczewski and Bowman 2020). Infection typically becomes visible within 12 to 15 days (Carson 1999; Hock et al. 1995; Kleczewski et al. 2019; Telenko et al. 2021) as brown-black, raised, semicircular fungal bodies (stromata) on the surfaces of corn foliage that resemble spots of tar. Stromata also may occur on stalks and ear leaf husks. In severe cases, leaf blights and early senescence may occur, leading to plant death (Carson 1999; Hock et al. 1995; Parbery 1963; Valle-Torres et al. 2020). Tar spot is characterized by signs and symptoms of small black fungal fruiting structures (stromata) scattered over the leaf surface and necrotic lesions that likely lead to reduced photosynthetic efficiency, resulting in poor grain fill, reduced silage quality, and poor yields. Corn yield losses of 40-60% can occur in severely affected fields in North America with some fields experiencing complete crop loss if tar spot is not managed (Hock et al. 1995; Mottaleb et al. 2019; Pereyda-Hernández et al. 2009; Telenko et al. 2020a). Tar spot signs and symptoms usually progress from the lower to the upper canopy under favorable conditions (Bajet et al. 1994; Hock 1989), but in the Midwest U.S. and Canada, “top-down” disease progression also has been observed, particularly when the pathogen moves into new areas (Valle-Torres et al. 2020).

The conditions that facilitated the emergence and rapid spread of *P. maydis* in the U.S. and Canada are unclear. Valle-Torres et al. (2020) and Broders et al. (2022) hypothesized that *P. maydis* may have emerged in northern North America due to an introduction from Puerto Rico, Mexico, or other Central American countries through several possible pathways, including the movement of infected plant materials by humans and possible long-distance dispersal of spores by wind, rain, or tropical storms such as hurricanes. Despite the persistence of *P. maydis* in cornfields across the U.S. and Canada, there is limited information on the pathogen’s genetic variation, gene flow, and population structure, which may influence disease severity and affect the efficacy of the host resistance component of disease management. One or multiple cycles of sexual reproduction annually may help generate genotypic diversity in the *P. maydis* population in the U.S. leading to high variation and diversity and consequent changes in its population genetic structure.

Gene flow is an evolutionary factor that substantially defines the population structure of many plant pathogen species (McDermott and McDonald 1993). Gene flow is the movement of genes into or out of a population (McDermott and McDonald 1993; Slatkin 1985). This can occur in fungi by the movement of spermatia, or migration of individuals and even a diverse species population. Gene flow leads to mixing of alleles, which results in genetic homogeneity among populations in the absence of natural selection and genetic drift. On the other hand, when gene flow is restricted population divergence via selection and genetic drift will not be mitigated, leading to genetic differentiation and eventually speciation (Heip et al. 1998). Its impact on the population structure is estimated based on the migration rate (Nm) of individuals, i.e., the number of individuals that would be exchanged between populations per generation to account for the observed population differentiation (Giraud et al. 2008). An essential factor to consider in devising management strategies against diseases is variation within pathogen populations (McDonald 1995; Grünwald et al. 2017). Detailed investigations of pathogen genetic diversity and population genetic structure in different geographical regions are required, as these reflect the history and the evolutionary potential of the pathogen (McDonald 1997). Furthermore, understanding the variation and genetic diversity within and among regional populations of a pathogen can help locate its possible center of origin (Stukenbrock and McDonald 2008). Information on the center of origin can likely dictate the pathogen’s ability to cause disease and its response to management.

Molecular tools have been helpful over the years in uncovering the genetic structure of many plant pathogen populations worldwide (Dutech et al. 2007; Gautier et al. 2014; Medini and Hamza 2008; Morgante and Olivier 1993). Some frequently used tools include Restriction Fragment Length Polymorphisms (RFLPs), Random Amplified Polymorphic DNA (RAPD), Amplified Fragment Length Polymorphisms (AFLPs), and Simple-Sequence Repeats (SSRs) or microsatellite markers. Of these molecular tools, SSR markers are highly favored for genetic analyses in population studies because they are species specific, multiallelic, reproducible, highly polymorphic, and are easily amplified via polymerase chain reactions (PCR) (Dutech et al. 2007; Gautier et al. 2014; Medini and Hamza 2008; Morgante and Olivieri 1993; Winter and Kahl 1995). SSR markers are cost effective and provide a more reliable interpretation of the population’s genetic diversity (Guichoux et al. 2011). They can help answer many questions in fungal population biology and genetics. For instance, they have helped understand the diversity levels, sources of variation, pathogen dispersal, reproductive mode, and host selection within several rust pathogen populations such as *Melampsora larici-populina, Puccinia graminis,* and *P. striiformis f.sp. tritici* (Ali et al. 2014; Barres et al. 2012; Berlin et al. 2012; Danies et al. 2014; Dutech et al. 2007).

The goal of this research was to develop a set of polymorphic microsatellite markers that would allow for effective genotyping of *P. maydis* isolates for genetic analyses of its populations within the U.S., and to use those markers to test the following four hypotheses: 1) that genetic differences exist among *P. maydis* isolates across corn-production areas in the U.S.; 2) that *P. maydis* populations in the Midwest U.S. are geographically structured; 3) that there is gene flow; and 4) that there is evidence for sexual recombination within *P. maydis* populations across corn-producing areas in five Midwestern states. Testing these hypotheses will provide a baseline about the genetic structure of *P. maydis* populations soon after its introduction into the U.S., its evolutionary potential, and can help inform the development of future management approaches.

## Materials and Methods

### Sample collection procedure

Corn leaf samples exhibiting tar spot symptoms were collected from fields in five affected states (Illinois, Indiana, Iowa, Michigan, and Wisconsin) in the Midwest U.S. from 2018 to 2020 (Table 1 and Figure 1). Samples from Indiana that were submitted to the Purdue University Plant and Pest Diagnostic Laboratory in 2015 and 2017 also were included. Additional collection details are presented in Supplementary Table 1. All samples were dried by pressing in newspaper and stored at room temperature until they were processed for DNA extraction.

**Table 1.** Sampling information of *Phyllachora maydis* from five Midwestern states.

| Region | Isolate ID | No. of samples | No. of counties | Collection year |
| --- | --- | --- | --- | --- |
| Iowa (IA) | IA01 - IA37 | 37 | 29 | 2019-2020 |
| Illinois (IL) | IL01 - IL19 | 19 | 18 | 2019-2020 |
| Indiana (IN) | IN01 - IN95 | 95 | 50 | 2015-2020 |
| Michigan (MI) | MI01 - MI10 | 10 | 6 | 2019-2020 |
| Wisconsin (WI) | WI01 - WI20 | 20 | 17 | 2019-2020 |
| Total |  | 181 | 120 |  |

**Figure 1.**
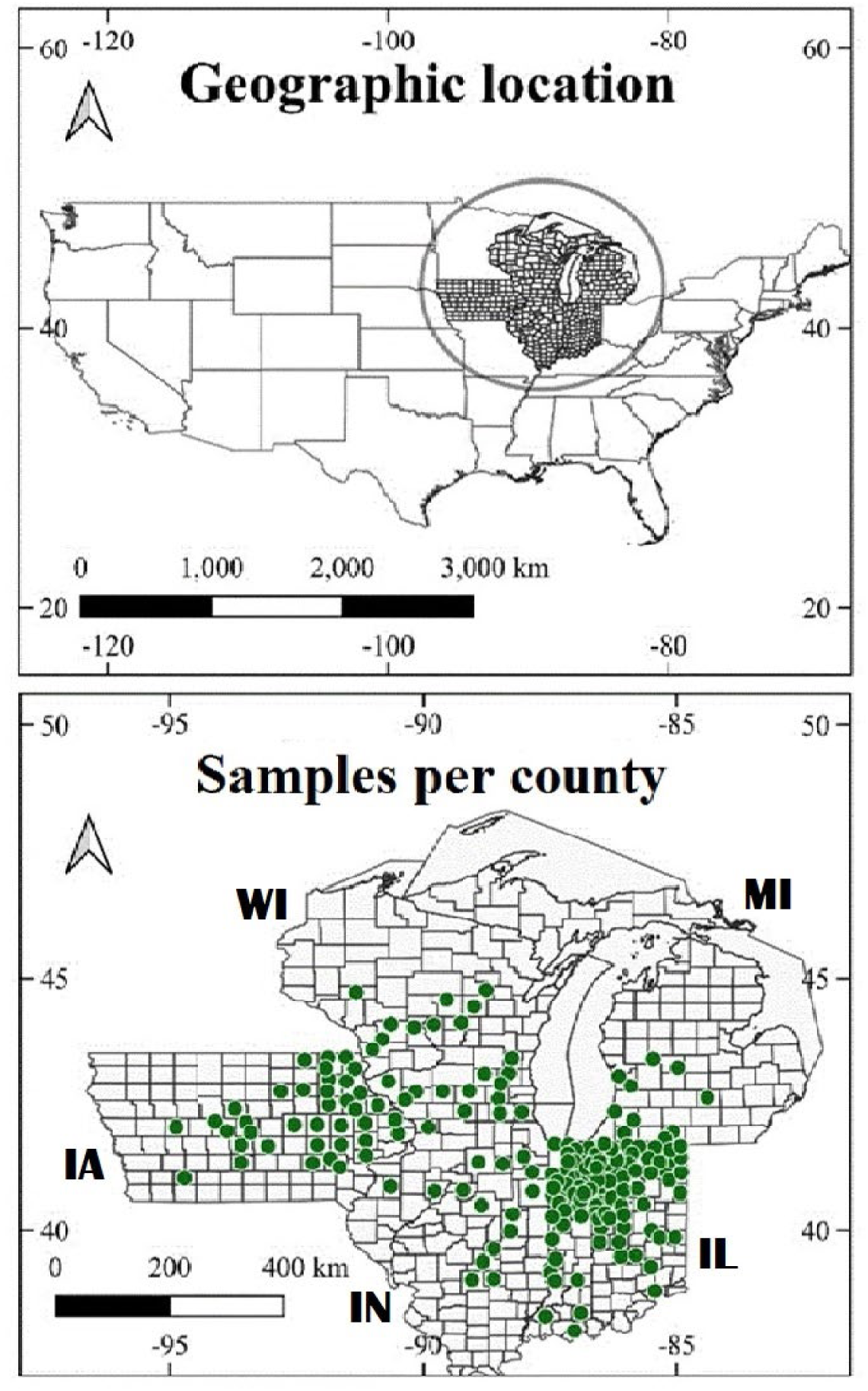
Geographic locations of the 181 *Phyllachora maydis* samples obtained from Midwest U.S. Green dots denotes number of samples analyzed per each county.

### DNA extraction and identification of fungal samples

Corn leaf samples approximately 8 cm in length with tar spot stromata were placed in a solution containing 2.5% (v/w) commercial bleach (a.i. 5.0% sodium hypochlorite) with one drop of Tween 20 added to every 50 ml of solution (Bio-Rad Laboratories, Hercules, CA) for 10 s to disinfest their surfaces. Samples were then rinsed twice with sterile water, placed on paper towels and air dried. Three to five stromata (> 5.2 mm in diameter) of *P. maydis* without necrotic halos and in close proximity to each other were excised from each leaf sample using a sterile surgical blade. Care was taken to limit the samples to stromata with the least amount of leaf material possible. The three to five excised stromata were pooled for each sample and placed in sterile 2-ml microcentrifuge tubes for processing. Each 2-ml microcentrifuge tube represented one *P. maydis* sample (isolate) for DNA extraction. The total genomic DNA (gDNA) from 181 *P. maydis* samples was recovered using a modified cetyltrimethylammonium bromide-based protocol (CTAB) (Healey et al. 2014). The DNA from each sample was measured with a Nanodrop 1000 spectrophotometer (Thermofisher Scientific, Waltham, MA), adjusted to a concentration of 0.2 ng/μL, and stored in nuclease-free sterile water at 4 °C until further molecular analyses. A *P. maydis* species-specific conventional PCR assay was used to confirm the identity of all samples before microsatellite analysis. This method can identify *P. maydis* by targeting part of the internal transcribed spacer (ITS1-5.8S-ITS2) region using primers ITS1F and ITS4 primers (Gardes and Bruns 1993; White et al. 1990). The amplified DNA sequences were analyzed by BLAST against a database generated from a partial assembly of the *P. maydis* genome (GenBank accession number: JAALGG000000000; Telenko et al. 2020b). The ITS region was located on a 0.418-Kb contig (GenBank accession number: ON241789). The sequence of this contig was used to identify a pair of species-specific primers (*P. maydis*-specific F and R) for the conventional PCR assay. The sequences of this pair of primers were then BLAST searched against The National Center for Biotechnology Information (NCBI) database to ensure they were specific to *P. maydis* sequence and were then synthesized by Integrated DNA Technologies (Coralville, IA). To test the specificity of the pair of synthesized *P. maydis* species-specific primers via conventional PCR assay, DNA extracted from ten *P. maydis* samples was amplified. Uninfected corn DNA and PCR mix without DNA were used as the controls in this assay to ensure specificity to *P. maydis*. Once specificity was confirmed, primers were then used to confirm *P. maydis* identity for the remaining samples used in this study.

### Microsatellite marker development and primer design

Primers for thirteen microsatellite loci (Table 2) were developed from the draft genome sequence data of *P. maydis* (GenBank accession number: JAALGG000000000). For SSR prediction, the *P. maydis* draft genome sequence was used as input in QDD version 3.1.2b (Meglécz et al. 2014). This program uses a set of Perl scripts that integrates the program Primer3 (Koressaar and Remm 2007) for designing primers flanking each SSR following the protocol outlined in Diaz-Valderrama and Aime (2016). The Primer3 parameters were as follows: primer lengths of 18 to 22 base pairs (bp); primer melting point (Tm) of 60 to 62 °C; and product sizes of 141 to 325 bp. Primer pairs for thirty microsatellite loci containing two- or three-nucleotide motifs were screened initially for polymorphism against a subset of ten isolates of *P. maydis* collected from five U.S. states during different years. Screening was repeated three times to ensure the microsatellites were reproducible. Among the thirty microsatellite primer pairs tested, thirteen were selected based on their ability to consistently amplify across all ten tested *P. maydis* samples in the three repeated screens and were polymorphic (Table 2).

**Table 2.**
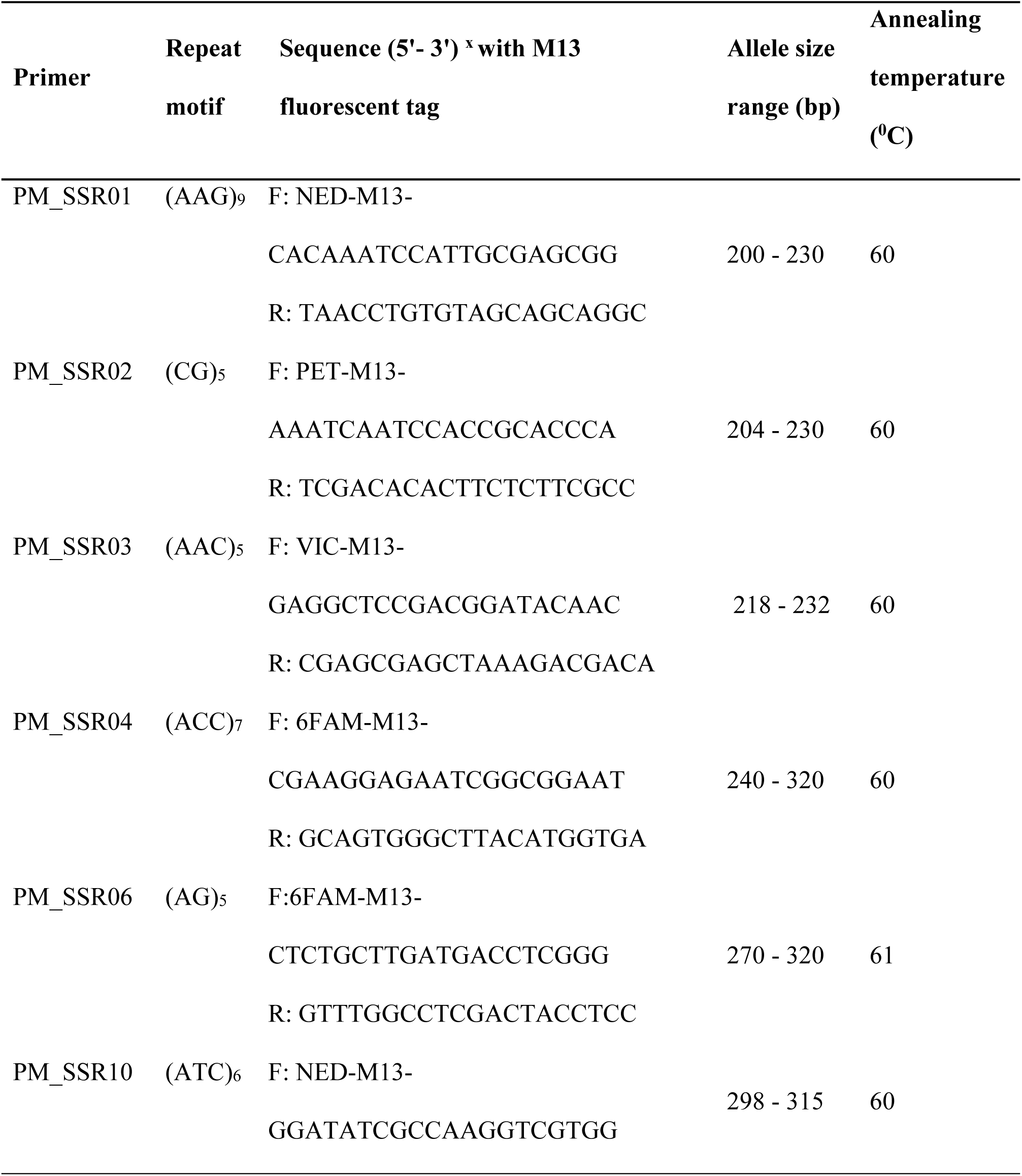
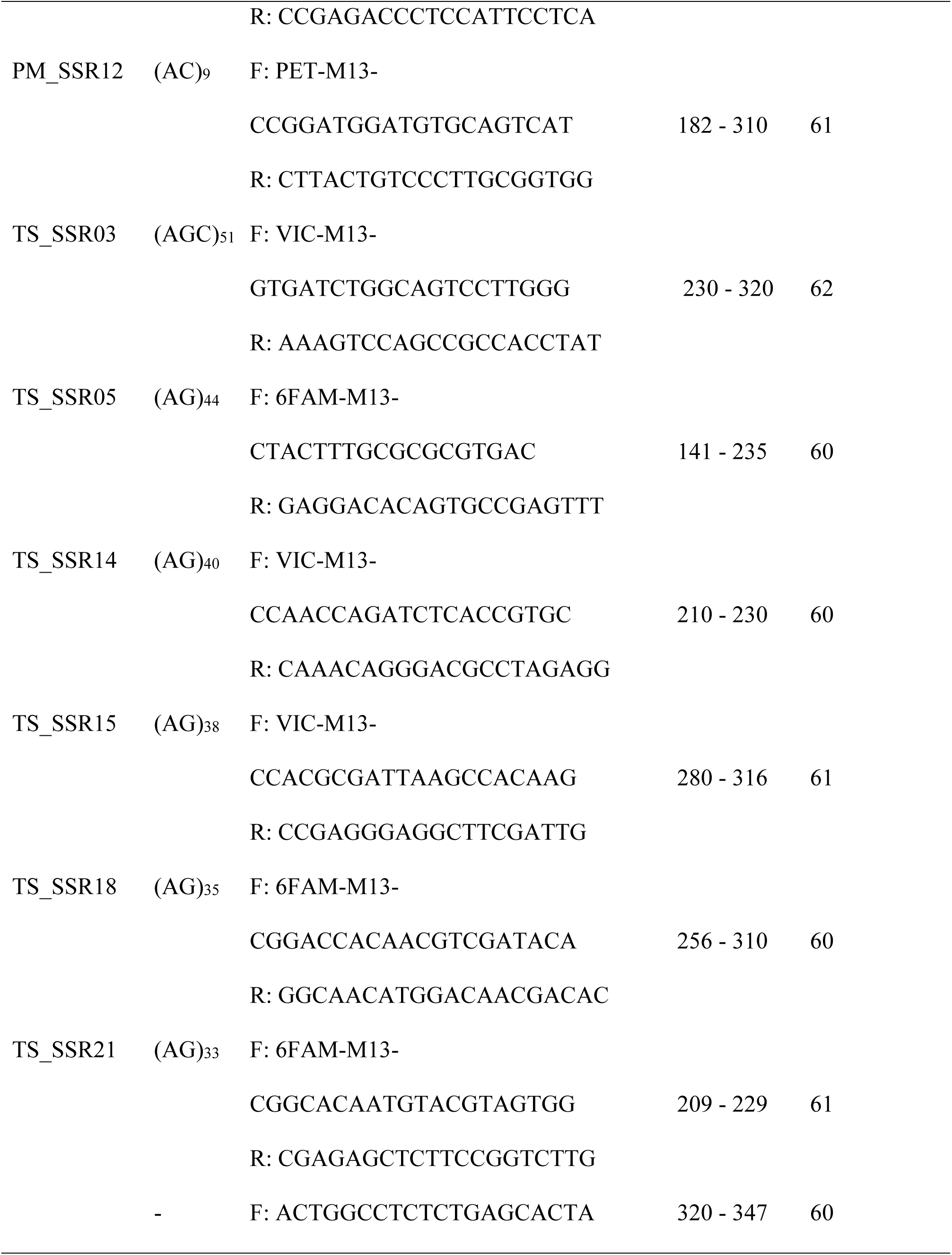
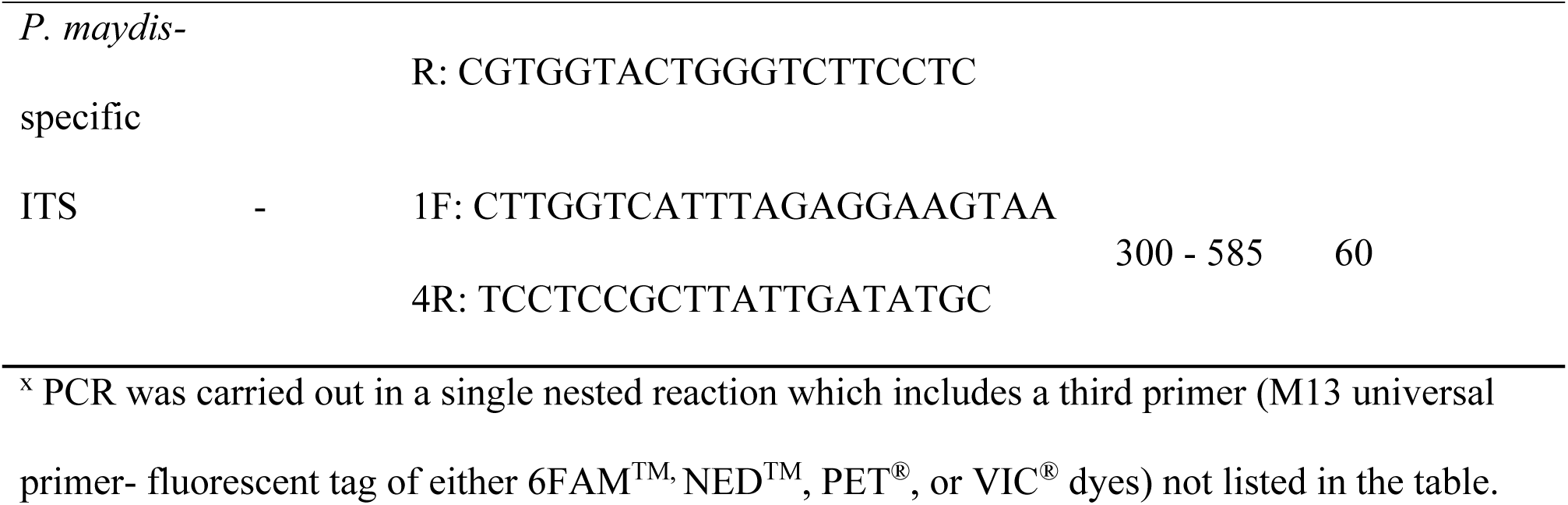
Sequences and properties of the 13 microsatellite loci used to analyze five populations of *Phyllachora maydis* in the Midwest U.S.

### SSR amplification and genotyping

Microsatellite loci were prepared for fragment analysis using a modified M13 method outlined by Schuelke (2000) and Diaz-Valderrama and Aime (2016). Each forward SSR primer contained an 18-bp tail added at its 5’ end with universal M13 primer (5’-TGTAAAACGACGGCCAGT-3’) which was previously labeled at its 5’ end with one of four specific fluorescent dyes (6-FAM^®^, NED^®^, PET^TM^ and VIC^TM^; Thermo Fisher Scientific, Waltham, MA, U.S.A) (Table 2). PCR amplifications were carried out in a final volume of 12.5 μL, containing 6.25 μL of Taq 2x Master Mix (New England Biolabs, Ipswich, MA, U.S.A), 0.16 μL of the forward primer with the M13 tail, 0.47 μL of the M13 primer with one of the four fluorescent dyes and 0.63 μL of the reverse primer (all at a concentration of 10 uM), and 5.0 μL of DNA template (0.2 ng/μL). Thermal cycling included 94 °C for 5 min; 30 cycles of 94 °C for 30 s, annealing temperature for microsatellite primer for 45 s, 72 °C for 45 s; eight cycles of 94 °C for 30 s, 53 °C for 45 s, 72 °C for 45 s; and 1 cycle of 72 °C for 10 min.

PCR products were pooled post PCR by physically mixing four aliquots each of PCR products labeled with different fluorescent dyes and sent to CD Genomics (Shirley, NY, U.S.A) for fragment analysis via capillary electrophoresis, which utilizes an ABI PRISM 3730xl automated sequencer. Fragment size analysis was returned, and allele size scoring of the fragments was performed using GeneMarker software 3.0.1 (SoftGenetics, State College, PA U.S.A.). Each SSR primer pair was presumed to amplify a single genetic locus and bands of different molecular weights were considered as different alleles. Amplification, sequencing, and genotyping of *P. maydis* samples were replicated three times with the same DNA preparations to ensure allele prediction due to the obligate nature of the pathogen where fungus co-exist with the host making it difficult to get completely pure DNA during extraction.

### Microsatellite diversity indices across populations

Microsatellite loci summary statistics were calculated using the ‘locus_table’ function within the R package poppr v 3.0.2 (Kamvar et al. 2014) and GenAlEx software v 6.501 (Peakall and Smouse 2012). Locus diversity indices included: the number of alleles (Na), Nei’s (1978) unbiased gene diversity (He), polymorphic information content (PIC) (Botstein et al. 1980), and genetic evenness (E_5_) which estimates the uniformity of genotype distribution across populations and ranges from 0 to 1. E_5_ = 1 means genotypes occur at equal frequency, regardless of richness (Grünwald et al. 2003).

### Population genetic diversity

Population genetic diversity analyses were estimated from the scored marker data. Genetic diversity measures the richness and abundance of genotypes in a population. It was estimated based on two indices: the Shannon-Weiner index of multilocus genotypic diversity (MGD),(I) (Shannon 2001) and Simpson’s complement index of MLG diversity (λ) (Simpson 1949) with the R package poppr v 3.0.2. Simpson’s (1949) index measures the probability that two individuals chosen at random from the population will be found to belong to the same group. Additionally, poppr v3.0.2, was used to identify unique MLGs and expected MLGs at the smallest sample size (eMLG) based on microsatellite allele sizes. The Na, Ne (number of effective alleles), Pa, E_5_, PIC, and He were estimated with poppr v 3.0.2 and GenAlEx v 6.501.

### Linkage disequilibrium

Linkage disequilibrium test (index of association, I_A_ and *r̄_<u>d</u>_*) (Agapow and Burt 2001) was done using the R package poppr v.3.0.2 with 999 permutations using the ‘ia’ function to determine if populations are clonal or sexual. In clonally reproducing populations, significant disequilibrium (α = 0.001) is expected due to linkage among loci whereas in sexually reproducing populations linkage among loci is not expected (Brown et al. 1980). The null hypothesis tested is that alleles observed at different loci are not linked if populations are sexual, while alleles recombine freely into new genotypes during sexual reproduction.

### Population genetic differentiation, gene flow, and structure analysis

Tests for population genetic differentiation (F_ST_ and *p* values) over 999 bootstrap replications were performed and gene flow (Nm) was estimated using the GenAlEx software. An analysis of molecular variance (AMOVA) and estimate of pairwise F statistics (F_ST_) among the groups were also performed to measure the probable differentiation among different groups and assessment of gene migration among populations over time (gene flow [Nm]) using GenAlEx software.

Population structure was further assessed with STRUCTURE version 2.3.4 (Pickard et al. 2000), based on the individual-based Bayesian clustering methods. A continuous series of K values from 1 to 10 was tested in 10 independent runs to deduce the optimal K value for the genotypes using the ΔK method (Evanno et al. 2005). Each run comprised a burn-in length of 100,000 followed by 100,000 MCMC (Markov Chain Monte Carlo) replicates. The most likely values of K were chosen based on ΔK that was computed with Structure Harvester version 0.6.94 (Earl and Von Holdt 2012). The optional alignment of clusters across individual runs for each K was determined using CLUMPP version 1.1.2 (Jakobsson and Rosenberg 2007), which included a greedy algorithm and 10,000 random input orders of 10 independent STRUCTURE runs.

The genetic-structure plot was drawn by Distruct version 1.1 software (Rosenberg 2010). The population structure based on the genetic distance among all sampled individuals was further revealed by principal coordinate analysis (PCoA) using GenAlEx software. Additionally, discriminant analysis of principal components (DAPC) was performed using the find.clusters command of the R package adegenet v 1.3 (Jombart et al. 2010) to identify the genetically differentiated clusters across the *P. maydis* studied populations.

Lastly, a genetic dissimilarity matrix was computed based on the continuous Euclidian dissimilarity index and Nei’s standard genetic distance (DST, corrected) (Nei 1972) over 1,000 bootstrapped replications based on the Unweighted Pair Group Method with Arithmetic Mean (UPGMA) trees were generated using ‘genpop’ with R package adegenet v1.3.

## Results

### Microsatellite polymorphism and gene diversity

A total of 181 *P. maydis* samples were genotyped using 13 polymorphic SSR markers (Table 3). Among the entire population 85 alleles were recovered and the number of alleles per locus (Na) varied from 3 to 13 with an average of 6.5. The highest number of alleles (Na = 13) was detected at locus TS_SSR15 (Table 3). For the entire studied population, gene diversity (He) (equivalent to the expected heterozygosity in a diploid) among loci ranged from 0.35 (PM_SSR06) to 0.76 (TS_SSR15) with an average of 0.55 (Table 3). Genetic evenness (E_5_) of markers ranged from 0.55 (PM_SSR06) to 0.88 (TS_SSR03) with an average of 0.70 (Table 2.3). Informativeness of individual loci as measured by their polymorphic information content (PIC) ranged from 0.32 (PM_SSR06) to a maximum of 0.72 (TS_SSR15) with a mean PIC value of 0.50 (Table 3).

**Table 3.** Summary statistics of 181 samples of *Phyllachora maydis* for each of the 13 analyzed microsatellite loci.

| Locus | Na<sup>z</sup> | He | E<sub>5</sub> | PIC |
| --- | --- | --- | --- | --- |
| PM_SSR01 | 7 | 0.73 | 0.79 | 0.69 |
| PM_SSR02 | 9 | 0.73 | 0.81 | 0.69 |
| PM_SSR03 | 5 | 0.50 | 0.75 | 0.42 |
| PM_SSR04 | 7 | 0.62 | 0.74 | 0.54 |
| PM_SSR06 | 4 | 0.35 | 0.55 | 0.32 |
| PM_SSR10 | 5 | 0.55 | 0.81 | 0.45 |
| PM_SSR12 | 6 | 0.52 | 0.60 | 0.47 |
| TS_SSR03 | 5 | 0.52 | 0.88 | 0.41 |
| TS_SSR05 | 3 | 0.49 | 0.72 | 0.43 |
| TS_SSR14 | 8 | 0.57 | 0.63 | 0.52 |
| TS_SSR15 | 13 | 0.76 | 0.65 | 0.72 |
| TS_SSR18 | 8 | 0.48 | 0.60 | 0.45 |
| TS_SSR21 | 5 | 0.38 | 0.54 | 0.35 |
| Mean | 6.5 | 0.55 | 0.70 | 0.50 |

<sup>z</sup>Na = number of observed alleles. He = Nei's (1978) unbiased gene diversity. E<sub>5</sub> = population evenness (estimates uniformity of genotype distribution) (Grünwald et al. 2003), PIC = Polymorphic information content (Botstein et al. 1980).

### Population genetic diversity

Table 4 summarizes the genetic diversity estimates across all populations averaged over the 13 microsatellite loci. All samples were unique and hence 181 unique multilocus genotypes were recovered from the 181 samples (Table 4). A genotype accumulation curve determined that 100% of the unique multilocus genotypes (MLGs) could be detected with eleven or twelve microsatellite markers (Figure 2). Genetic diversity estimates across populations had a mean number of alleles (Na) of 3.7 with a range of 2.7 to 5.5, the mean number of effective alleles was 2.1 (range 1.9 to 2.4), and the frequency of private alleles (Pa) or those that were unique to a single population was 0.45 (range 0.15 to 1.31) (Table 4). All indices of genotypic diversity were high: Shannon-Weiner’s index (I), a measure of population richness (biodiversity), ranged from 0.82 to 1.03 with a mean diversity index of 0.84; and Simpson’s complement index (λ) ranged from 0.90 to 0.99 with mean index of 0.99 (Table 4). Likewise, Nei’s unbiased gene diversity (He) was high for all populations and ranged from 0.41 to 0.53 (Table 4). The genotypic diversity (E_5_ = 1) was evenly distributed in all populations (Table 4). The Indiana population had the highest Na, Ne, Pa, I, λ, and He values, whereas the Michigan population had the lowest values for all statistics (Table 4). The average percentage of polymorphic loci (PPL) per population was 94% with a range from 85% (Michigan) to 100% (Illinois and Indiana). The PPL for both Iowa and Wisconsin were 92% (Table 4).

**Figure 2.**
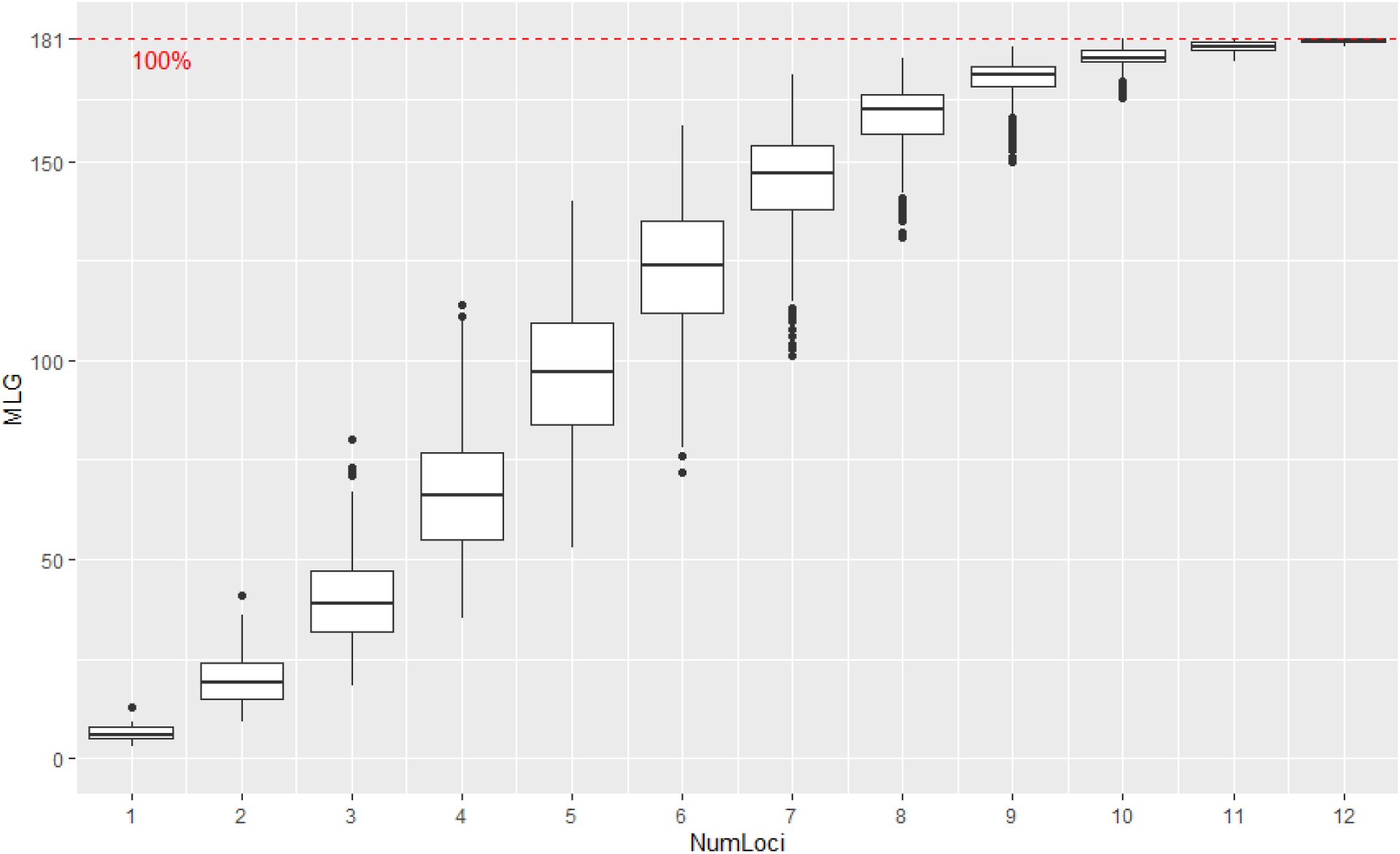
Genotype accumulation curve for unique multilocus genotypes (MLG) of *Phyllachora maydis* detected as the number of microsatellite loci (NumLoci) sampled. When 11 or 12 microsatellite loci are used in genotyping, 100% of the unique multilocus genotypes can be detected. The 100% unique multilocus genotype level is indicated by the red dashed line.

**Table 4.**
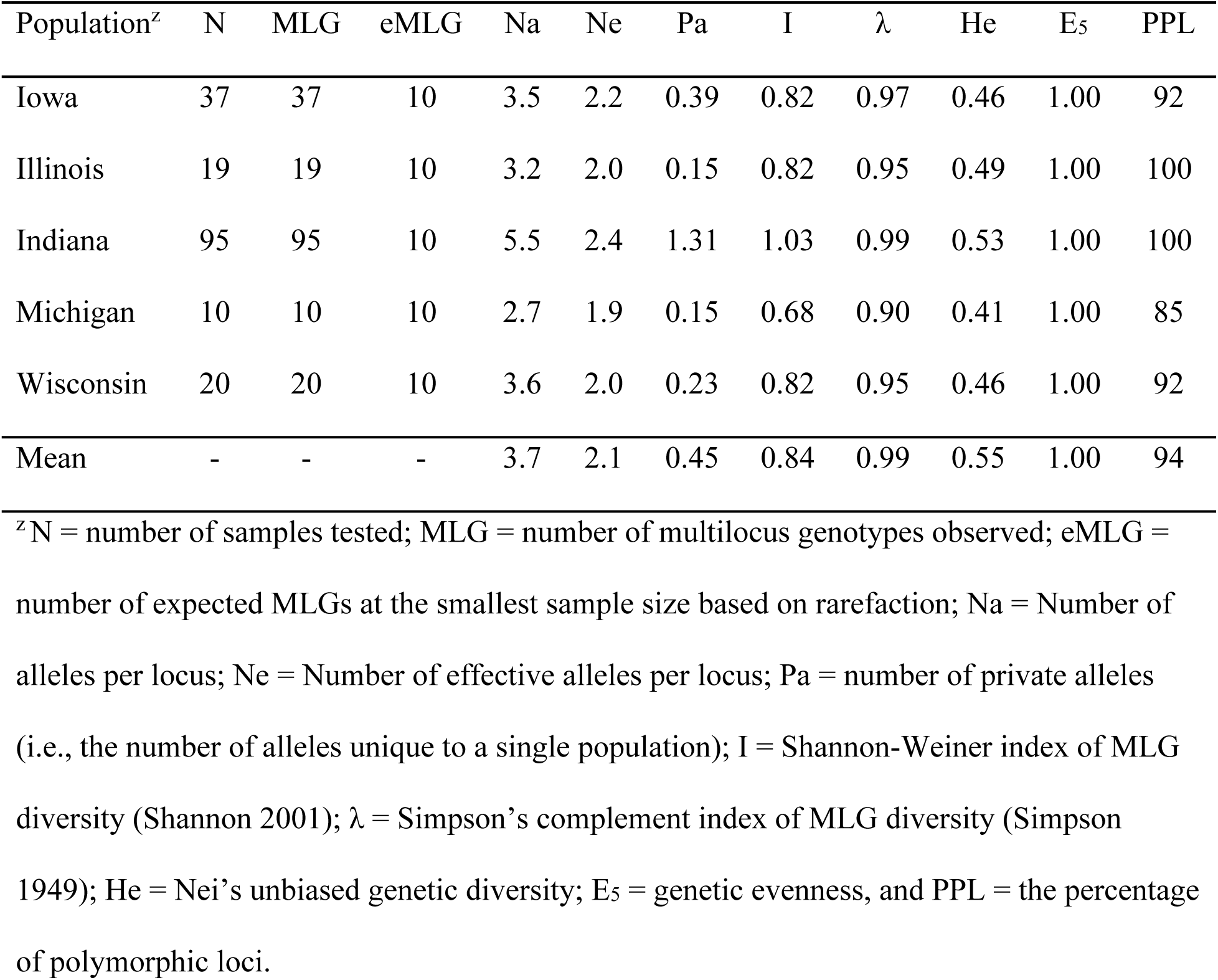
Genetic diversity indices across all populations averaged over the 13 SSR loci.

### Linkage disequilibrium (LD)

The test for linkage disequilibrium (LD) using the microsatellite data of the 181 samples by the index of association (I_A_) and the standard index of association (*r̄_<u>d</u>_*) using 999 permutations showed evidence that all populations are likely reproducing sexually (Table 5). The linkage disequilibrium null hypothesis of no linkage among loci failed to be rejected since in all populations *P* > 0.001 suggesting that alleles are not linked, and populations are likely sexual (Table 5).

**Table 5.**
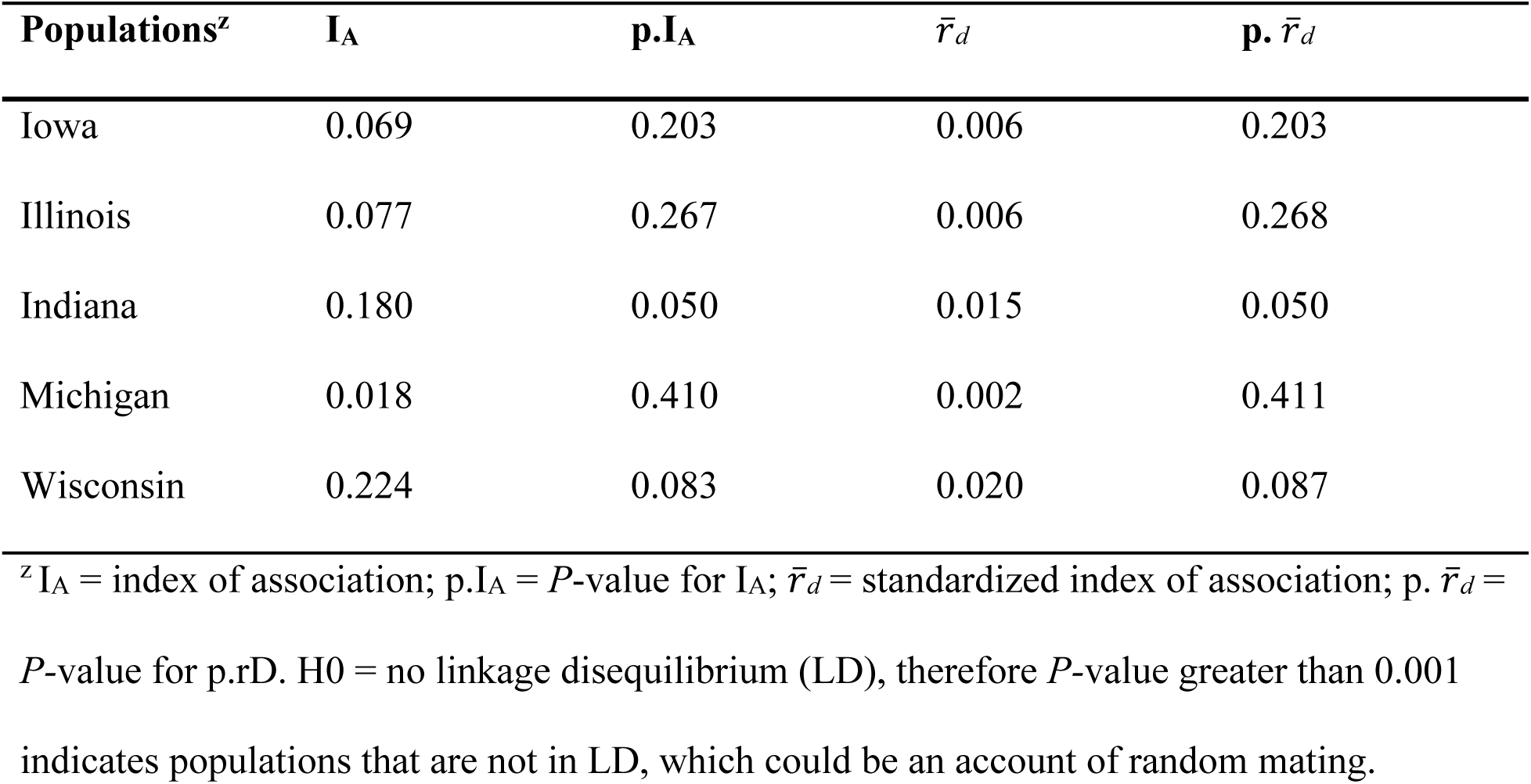
Linkage disequilibrium (LD) based on index of association test (I_A_ and *r̄_<u>d</u>_*) across the five regional populations.

| Populations<sup>z</sup> | $I_A$ | $p.I_A$ | $\bar{r}_d$ | $p. \bar{r}_d$ |
| --- | --- | --- | --- | --- |
| Iowa | 0.069 | 0.203 | 0.006 | 0.203 |
| Illinois | 0.077 | 0.267 | 0.006 | 0.268 |
| Indiana | 0.180 | 0.050 | 0.015 | 0.050 |
| Michigan | 0.018 | 0.410 | 0.002 | 0.411 |
| Wisconsin | 0.224 | 0.083 | 0.020 | 0.087 |

<sup>z</sup> $I_A$ = index of association; $p.I_A$ = $P$ -value for $I_A$ ; $\bar{r}_d$ = standardized index of association; $p. \bar{r}_d$ =
$P$ -value for $p.rD$ . $H_0$ = no linkage disequilibrium (LD), therefore $P$ -value greater than 0.001
indicates populations that are not in LD, which could be an account of random mating.

### Population genetic differentiation, gene flow, and structure analysis

Analysis of molecular variance (AMOVA) based on the F statistics was estimated with and without grouping populations according to their geographical locations. AMOVA showed that 85% of the total variation was due to within-population variation and only 15% was accounted for by genetic divergence among populations (Table 6). This result was further validated by the principal coordinate analysis (PCoA) (Figure 3), where the first two axes explained 19.6% of the total variation, with each of the coordinates (1 and 2) accounting for 11.7% and7.9%, of the variation, respectively.

**Figure 3.**
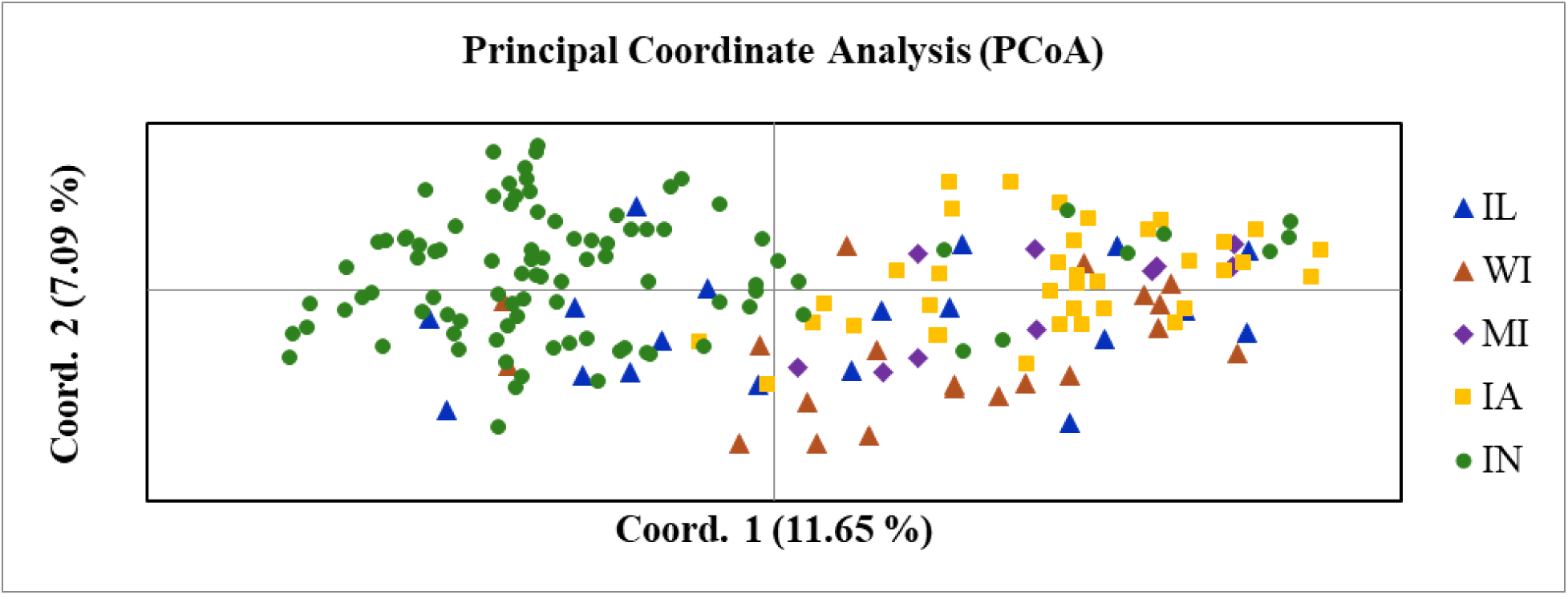
Principal coordinate analysis (PCoA) showing the clustering of 181 *Phyllachora maydis* samples as revealed by 13 microsatellite loci. Samples coded with the same color and shape belong to the same population. PCoA explained 19.6% of the total variation and the first two axes (1 and 2) accounted for 11.7% and 7.9% respectively. Population abbreviations are IA = Iowa, IL = Illinois, IN = Indiana, MI = Michigan, and WI = Wisconsin.

**Table 6.** Analysis of molecular variance (AMOVA) showing genetic variation within and among *Phyllachora maydis* populations in the Midwest U.S.

| Source | Df | SS | MS | Est. Var. | % Var. | F<sub>ST</sub> | Nm |
| --- | --- | --- | --- | --- | --- | --- | --- |
|  |  |  |  |  |  | (P-value) | Average |
| Among Populations | 4 | 157.36 | 39.34 | 0.61 | 15 | 0.15<br>(0.001) | 2.80 |
| Within Populations | 357 | 1221.56 | 3.42 | 3.42 | 85 |  |  |
| Total | 361 | 1378.93 |  | 4.03 | 100 |  |  |

Df = Degrees of freedom; SS = Sum of squares; MS = Mean sum of squares; Est. Var = Estimated variance; % Var. = percentage of variance, F<sub>ST</sub> = genetic differentiation statistic (Wright 1965), Nm = gene flow (Giraud et al. 2008).

The overall genetic differentiation among populations (F_ST_ = 0.15, *P* = 0.001) was relatively low. Similarly, pairwise F_ST_ values of genetic distances among all populations were statistically significant (*P* = 0.001) (Table 7). Among all populations, the average estimated gene flow, Nm, was 2.80 (Table 6). Discriminant analysis of principal components showed that *P. maydis* populations in the Midwest are not geographically structured based on collection origins (regions), but instead, saw an intermixing of *P. maydis* samples from Illinois, Indiana, and Wisconsin and then another intermixing of samples from Iowa and Michigan (Figure 4). A cluster dendrogram generated using the Unweighted Pair Group Method with Arithmetic mean based on nei_distance bootstrapped grouped the five regional populations into two clusters (1 and 2) that did not correspond with geography (Figure 5). Cluster C1 was composed of populations Illinois, Indiana, and Wisconsin and cluster C2 was composed of populations Iowa and Michigan (Figure 5).

**Table 7.** Population genetic differentiation measured by F_ST_ (below the diagonal) in pairwise comparisons among the five populations of *Phyllachora maydis* with p values (above the diagonal).

| $F_{ST}/P\text{-value}$ | IL | WI | MI | IA | IN |
| --- | --- | --- | --- | --- | --- |
| IL | --- | 0.001 | 0.001 | 0.001 | 0.001 |
| WI | 0.08 | --- | 0.001 | 0.001 | 0.001 |
| MI | 0.18 | 0.12 | --- | 0.001 | 0.001 |
| IA | 0.16 | 0.12 | 0.12 | --- | 0.001 |
| IN | 0.13 | 0.15 | 0.17 | 0.16 | --- |

IA = Iowa, IL = Illinois, IN = Indiana, MI = Michigan and WI = Wisconsin.

**Figure 4.**
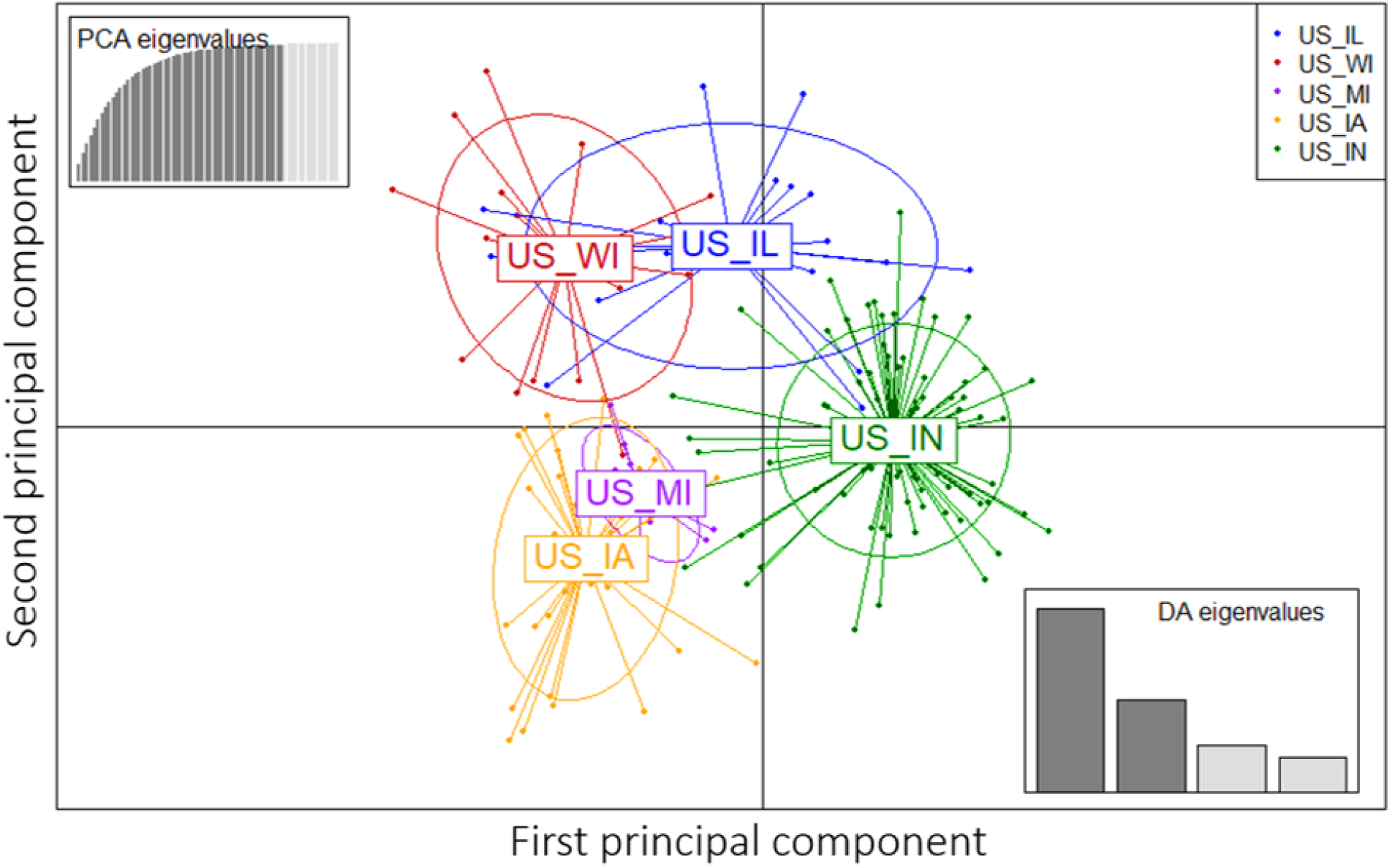
Discriminant analysis of principal component of five *Phyllachora maydis* populations in the Midwest. *Phyllachora maydis* (N =181) samples from the five populations in the Midwest U.S formed clusters of severe admixtures. Each color represents a population: yellow = Iowa (US_IA); blue = Illinois (US_IL); green = Indiana (IN); purple = Michigan (US_MI) and orange = Wisconsin (US_WI).

**Figure 5.**
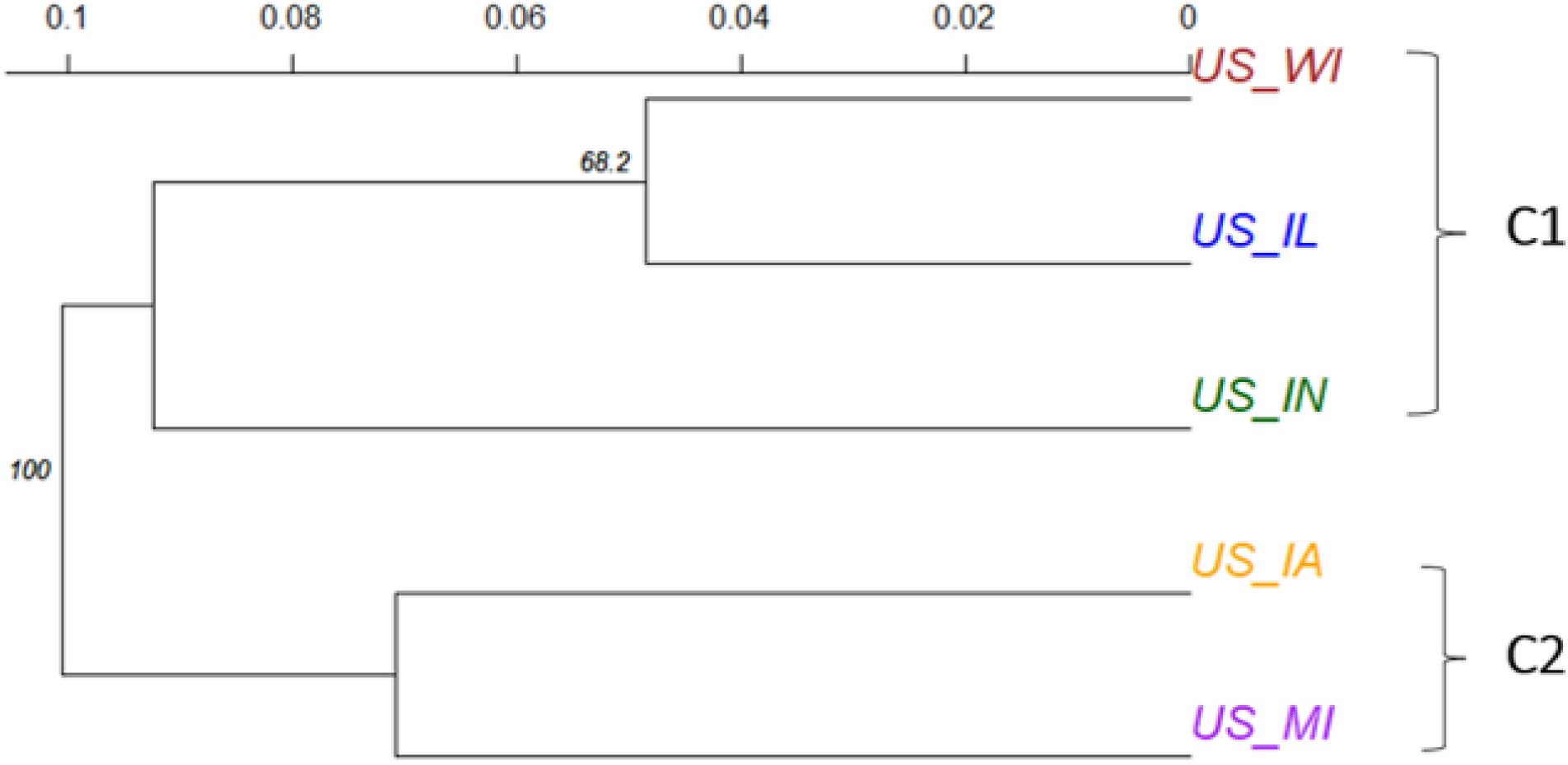
Dendrogram generated with the Unweighted pair-group with arithmetic mean method showing the genetic relationships among the five populations of *Phyllachora maydis* from the Midwest based on Nei’s (1972) genetic distance over 1,000 bootstrap replicates. *P. maydis* samples formed two clusters, C1 (Illinois, Indiana, and Wisconsin) and C2 (Iowa and Michigan). Numbers above branches represent percentage of bootstrap values, and values less than 60% were not indicated.

Finally, Bayesian clustering of the 181 isolates reached a sharp peak at K = 2 (Figure 6a), confirming that the 181 samples evaluated in this study could be most likely clustered into two genetic clusters. The result (bar plot) also detected a greater degree of genetic admixture between the two genetic clusters with no clear geographic origin-based structuring among the five geographical regions from which *P. maydis* was collected in the Midwest (Figure 6b). Similar genetic structure among the individuals which supported severe admixture within the two clusters was also confirmed by DAPC analysis.

**Figure 6.**
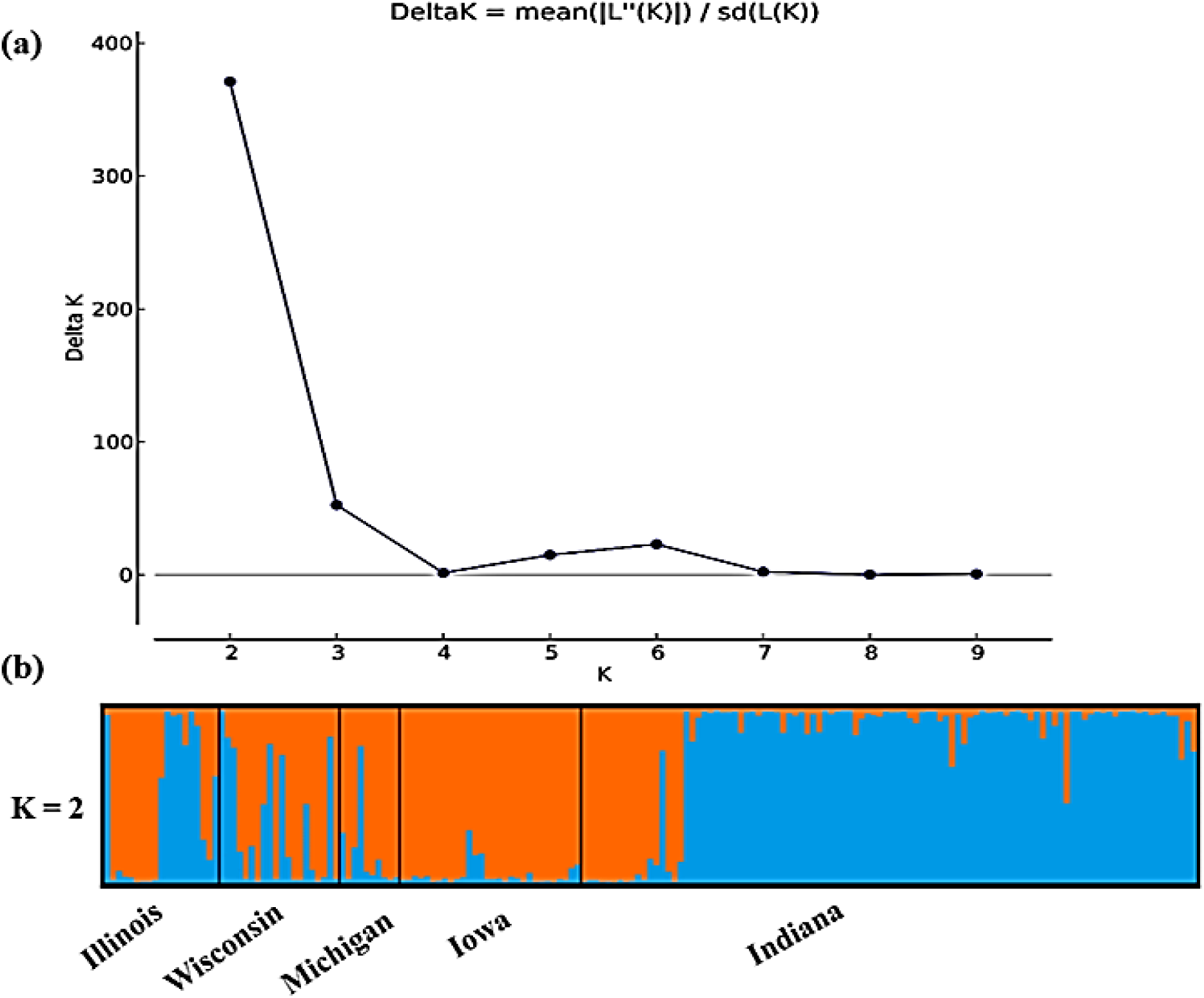
Population structure of 181 samples of *Phyllachora maydis* representing five populations in the Midwest U.S. **(a)** Best delta K value with maximum peak estimated using the method of Evano et al. (2005); and **(b)** Graph of the estimated population structure for K = 2 designated by Structure Harvester. The different colors (orange and blue) represent genetic groups or clusters. The x axis represents the origin of the samples (assigned populations), and the colored columns represent each isolate of *P. maydis* fragmented in K colored sections. The y axis represents the proportion of ancestry to each cluster.

## Discussion

This is the first analysis assessing the population dynamics of *Phyllachora maydis* populations using SSR markers. Knowledge on the population structure, genetic variation, diversity, mode of reproduction, and gene flow are essential to understanding how pathogen populations spread and overcome host resistance (McDonald and Linde 2002; Morris et al. 2014). Rampersad et al. (2013) emphasized that areas of high gene diversity may serve as sources for the emergence of new genotypes with novel biological characteristics and consequent changes in pathogen resistance to chemical compounds and increased fitness within their populations.

In this analysis, 13 highly polymorphic SSR markers were developed and used to assess the population dynamics of *P. maydis* samples from five corn-growing states in the Midwest U.S. The polymorphisms detected by these SSR markers ranged from reasonably informative (>0.25 PIC <0.50) to highly informative (PIC >0.50) (Reyes-Valdés 2013; Rosenberg et al. 2003). These SSR loci displayed allelic diversity among *P. maydis* samples, from 3 to 13 alleles per locus, where the PIC values ranged from 0.32 to 0.72. The PIC provides an estimate of the discriminatory power of a locus by considering the number and relative frequencies of the alleles (Marulanda et al. 2014). All 13 loci displayed differences for numbers of effective and private alleles, or those unique to a specific geographical area and are useful in comparing diversity between species or populations (Mahmodi et al. 2014). The Indiana population contained the highest number of private alleles compared to samples from the other states (Table 4). Although this is due at least in part to its larger sample size, the very large difference between the mean number of private alleles per locus in Indiana (1.31) versus the highest in any other state of only 0.39 in Iowa may reflect a biological difference. Possible explanations for this difference may be that the Indiana population received higher diversity from a possible donor population, is being a continuous receptor of *P. maydis* populations from an introduction/establishment pathway, and/or that is has existed longer so has had more time to accumulate new alleles. However, these would require a comparative analysis of *P. maydis* to the other species of *Phyllachora* to rule out the possibility of a host shift and allele accumulation over time. Additionally, since *P. maydis* most likely was introduced from source(s) outside the U.S., a quantitative introduction pathway risk analysis is required. Such assessment could benefit in the understanding of possible routes of entry and establishment of *P. maydis* in the U.S. For example, we could estimate the probable risks of the different introduction routes (by either imports of goods or weather events), establishment, and outbreak events (Cruz 2013). Additional sampling from other locations is required to test these hypotheses.

Results from AMOVA indicated that the highest percentage of variation (85%) was within populations of *P. maydis* and that gene diversity within the Midwest populations was high (He = 0.55) (Table 3). This high within-population diversity may be attributed to the recent emergence or introduction of the fungus to northern North America. Higher genetic diversity was observed in the Indiana (He = 0.53) and Illinois (He = 0.49) populations compared to those from other states, possibly due to the initial introduction and therefore detection of *P. maydis* in the U.S. in Indiana and Illinois during 2015. From a quantitative introduction pathway point of view, results might indicate that these states might be significant receptors (founder or potentially new arrivals). However, more research is needed to address this hypothesis. Nevertheless, the higher diversity in the Indiana population, compared to those in the other regions, may also be attributed to the larger number of assessed samples, thus revealing more genetic information of the population or the possibility of ongoing arrivals of the pathogen in the U.S. Lower genetic diversities were observed for the Iowa (He = 0.46), Wisconsin (He = 0.46), and Michigan (He = 0.41) populations, which could be associated with the later introduction or emergence of the pathogen into these areas.

The amount of gene diversity within a population may also be a function of population size where older populations have maintained higher levels of gene diversity compared to a recently colonized habitat (Rampersad et al. 2013). Differences in diversity levels across *P. maydis* populations in the Midwest U.S. may also be due to environmental conditions, geography, or corn genetics. Bennett et al. (2005) and Marulanda et al. (2014), both indicated that gene flow, sexual and asexual recombination could generate and affect genetic diversity within pathogen populations. This study shows evidence for that the regional populations are possibly sexual which was indicative by the based on the lack of linkage disequilibrium among loci (I_A_ and *r̄_<u>d</u>_*: *P* < 0.001). New genotypes can emerge during sexual reproduction and pathogens with active sexual cycles pose more significant risks (McDonald and Linde 2002). The existence of sexual recombination may give rise to new genotypes with advantageous allele combinations, which are potentially more adaptable to overcome host resistance, thus spreading diversity across populations for generations (Milgroom 1996). Nevertheless, clonally reproducing fungi may show as many alleles as those that undergo recombination, suggesting that gene diversity may not always be influenced by the reproduction mode of pathogens (McDonald 1997). Larger samples from the other states are needed to distinguish among these possibilities and to test whether the higher diversities in Indiana and Illinois reflect current and ongoing potential introductions or are due to sampling effects.

Analyses of *P. maydis* populations in the Midwest U.S. showed that *P. maydis* samples from the five sampled states were related or at least genetically alike based on their genetic identity and there was low but statistically significant genetic differentiation (F_ST_ = 0.15, *P* = 0.001) among Midwest populations. Further, results from principal coordinate, discriminate component and cluster analyses showed that *P. maydis* populations within the Midwest are not geographically structured; instead, populations had a greater degree of genetic admixtures with moderate gene flow (Nm = 2.80), also referred to as the migration rate, between *P. maydis* populations in this analysis. The relatively low degree of genetic differentiation and moderate gene flow observed among the *P. maydis* populations examined in this analysis may be due to inoculum dispersal over long distances even from locations outside of the U.S. This may have allowed the pathogen to spread among corn production areas in the Midwest U.S. since ascospores of *P. maydis* can be disseminated up to 1,200 m from the inoculum source (Kleczewski and Bowman 2020).

The exact mode of dissemination of *P. maydis* throughout the Midwest U.S. Corn Belt is not fully understood. However, the two most likely hypotheses include long-distance aerial dissemination of spores, or movement of infested corn material from one location to the next, presumably through human activities but also possibly via climatic systems such as high wind events including derechos and tornadoes (Broders et al. 2022; Valle-Torres et al. 2020). Dissemination of infested corn material may have resulted in migration and gene flow between populations resulting in the genetic signal of admixture among samples from the different geographical origins. Gene flow is promoted by several activities such as movement or exchange of infested plant material, and long-distance dispersal of spores (McDermott and McDonald 1993; McDonald 1997; McDonald and Linde 2002; Milgroom 1996; Milgroom and Peever 2003). Kawecki and Elbert (2004) indicated that divergence between pathogen populations could occur within local environments because of adaptation and genetic drift that increase relative fitness in different niches. When Nm > 1, migration will be sufficient to reduce genetic differentiation among populations (McDermott and McDonald 1993; Wright 1951).

Because *P. maydis* was not detected in the U.S. prior to 2015 (Ruhl et al. 2016), all populations in the Midwest presumably derive from one or more recent, possibly very limited, introductions. Therefore, our initial expectation was that genetic diversity within this area might be very low. For this reason, our strategy was designed to analyze only one or two samples per field across as wide an area as possible to capture the total level of genetic diversity. Grouping samples by state is an artificial construct to provide an initial assessment of regional genetic diversity that may not reflect biological realities. Analyses of better-defined populations can provide more accurate estimates of within- and among-population genetic diversity in the future. In the meantime, STRUCTURE analysis (Evanno et al. 2005) can provide a good initial indication of how the current genetic diversity is distributed.

STRUCTURE and population genetic analyses based on Nei’s (1972) genetic distance supported the subdivision of the *P. maydis* population into two clusters, suggesting the possibility of differentiation within the species. These results support the findings of Broders et al. (2022) that revealed variation and possibly multiple species related to *P. maydis* in northern North America. Identification of population subdivision within a particular geographic area could be associated with variations in the agro-ecosystems, such as sources of inoculum and host or tissue specificity (Milgroom and Peever 2003). The present analysis of *P. maydis* samples from five corn-producing states in the Midwest U.S. revealed higher-than-expected genetic diversity for a founder population and small but significant genetic differentiation among populations when assessed using 13 newly developed SSR markers. Analyses of additional samples with these molecular markers are needed to further assess populations from other affected regions within and outside the U.S. to gain a better understanding of the population genetics and dynamics of *P. maydis* in its source and founder populations.

The 13 highly polymorphic SSR markers identified in this study helped in understanding the current population structure, genetic variation, genetic diversity, reproductive mode, and gene flow of *P. maydis* in the Midwest U.S. The observed genetic diversity in all five populations was moderate to high and genetic variation among populations was low but significant, likely suggesting inter-state dispersal of sexually produced inoculum. The information generated in this study could be useful in understanding the biology and dissemination of *P. maydis* in the Midwest. This information provides a background that could aid in future studies on disease epidemiology, host-pathogen interactions, and help guide the development of disease management strategies. These 13 SSR markers could be useful for characterizing *P. maydis* samples from within the U.S. and other countries. The knowledge resulting from their use could further be useful in taxonomic and phylogenetic studies, functional genomics, genome mapping, gene tagging and quantitative trait linkage (QTL) analysis, hybrid testing and hybridization, and marker assisted-selection (MAS) breeding studies, pertaining to this pathogen and disease.

## Acknowledgements

The authors thank G. Ruhl, C. Da Silva, and S. Shim at Purdue University for assistance with sample collection and processing. The authors also thank C. Groves, B. Mueller, and B. Hudelson for collecting and curating samples at University of Wisconsin-Madison; and E. Roggenkamp, J. Byrne, J. Check, and A. McCoy at Michigan State University.

